# Antagonist binding actively disrupts interleukin-1 receptor dynamics to block co-receptor recruitment

**DOI:** 10.64898/2026.01.27.701974

**Authors:** Chandran Nithin, Ayomide Fasemire, Sebastian Kmiecik

## Abstract

The interleukin-1 receptor type 1 (IL1R1) is a central regulator of inflammatory signaling and functions as a molecular switch, yet it remains unclear how agonists and antagonists that bind the same primary site produce opposite signaling outcomes. Available structural data define inactive and active endpoint conformations but do not explain how antagonist binding dynamically prevents co-receptor recruitment.

Here, we combine all-atom molecular dynamics simulations with multiscale flexibility modeling using CABS-flex to systematically compare the intrinsic dynamics of IL1R1 across its unbound, agonist-bound, antagonist-bound, and co-receptor–bound states. Although both agonists and antagonists engage the same conserved interface on the D1/D2 domains, they induce fundamentally different dynamic responses in the receptor. Agonist binding progressively stabilizes interdomain coupling and promotes a stepwise transition toward a signaling-competent conformation. In contrast, antagonist binding selectively increases flexibility of the distal D3 domain, particularly at the co-receptor binding interface, thereby preventing progression along the activation pathway.

Importantly, the interaction patterns and dynamic signatures observed in the simulations are consistent with experimentally identified binding determinants, mutational data, and structural features associated with receptor activation and inhibition. These results demonstrate that IL1R1 antagonism is an active, allosteric, dynamics-driven process rather than a simple failure to stabilize an active conformation. Together, this work provides a mechanistic framework that reconciles existing structural and functional observations and highlights receptor dynamics as a key determinant of signaling control in the interleukin-1 system.

## 1.0 Introduction

The interleukin-1 (IL1) signaling system is a central regulator of innate immunity, and its dysregulation contributes to a wide range of inflammatory diseases, including rheumatoid arthritis, atherosclerosis, and cancer [1–6]. Therapeutic strategies targeting individual IL1 family cytokines often show limited efficacy due to functional redundancy within the family, where inhibition of one cytokine can be compensated by others [7,8].

Recent experimental studies have shown that multiple IL1 family cytokines, including IL1, IL-33, and IL-36, signal through a shared co-receptor, IL1 receptor accessory protein (IL1-RAcP), establishing it as a central hub for inflammatory signal integration [9]. This organization reflects a broader principle in cytokine signaling, in which multi-component receptor architecture and assembly, rather than ligand identity alone, determine signaling outcomes [10].

IL1 signaling operates as a molecular switch. The binding of an agonist cytokine to the primary receptor IL1R1 promotes recruitment of IL1-RAcP and formation of an active ternary signaling complex [11,12]. In contrast, the natural antagonist IL1-Ra binds the same receptor site with high affinity but fails to recruit the co-receptor, thereby preventing signaling. This functional dichotomy is encoded in the modular extracellular architecture of IL1-R1, in which the D1 and D2 domains mediate ligand binding, while the C-terminal D3 domain engages IL1-RAcP [13–17].

Structural studies have provided key snapshots of these functional states, summarized in Figure 1. In the antagonist-bound complex, the D1/D2 domains are clamped onto IL1-Ra while the D3 domain undergoes a pronounced reorientation that is incompatible with IL1-RAcP recruitment (Figure 1A, B) [13]. In contrast, agonist binding preserves a D3 orientation competent for co-receptor association, forming a primed signaling state (Figure 1C). Recent cryo-EM structures of the fully assembled ternary complex [18] further define the precise D3–IL1-RAcP interface required for receptor activation (Figure 1D).

**Figure 1.**
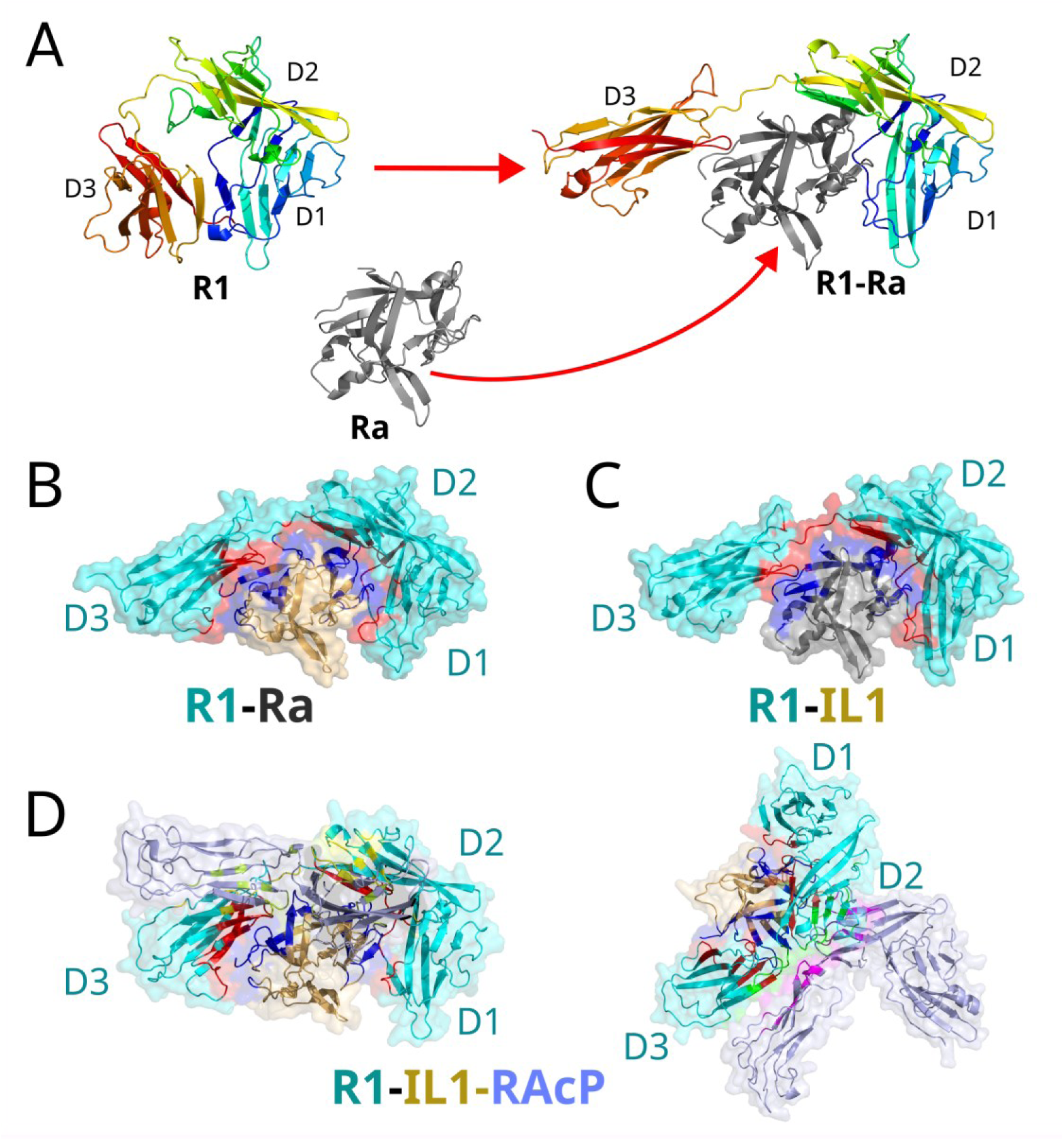
Structural basis of agonist and antagonist control of IL1 receptor signaling. (A) Binding of the antagonist Ra induces a major reorientation of the D3 domain relative to the unbound R1 receptor, resulting in an inactive receptor conformation. (B) In the antagonist-bound complex (R1–Ra; PDB: 1IRA), the altered positioning of the D3 domain prevents recruitment of the co-receptor RAcP. (C) Agonist binding (IL1) to R1 in the absence of the co-receptor (R1–IL1) preserves a D3 orientation compatible with subsequent RAcP recruitment, forming a primed signaling state (PDB: 1ITB). (D) Recruitment of RAcP to the primed R1–IL1 complex yields the fully assembled, signaling-competent ternary complex (R1–IL1–RAcP; PDB: 4DEP). Two alternative orientations of the ternary complex are shown to clearly visualize the position of the co-receptor RAcP relative to R1 and the agonist IL1. In panels B–D, the primary ligand-binding interface on R1 is highlighted in red, while the corresponding interface on Ra or IL1 is shown in dark blue. The secondary co-receptor binding interface is highlighted in yellow on R1 and green on the co-receptor RAcP.

Together, these observations establish the D3 domain as a critical checkpoint for signaling competence, a conclusion reinforced by therapeutic antibodies that block IL1-RAcP and suppress signaling from multiple IL1 family cytokines [9,19]. Collectively, these findings indicate that the geometry and organization of the D3–IL1-RAcP interface, rather than ligand affinity alone, determine signaling outcomes.

Despite this progress, static structures capture only end states of the signaling process and do not explain how the receptor transitions between inactive, primed, and active conformations. Key mechanistic questions therefore remain unresolved: how flexible the unbound receptor is, how do agonists and antagonists differentially reshape receptor dynamics, and why antagonist binding disrupts co-receptor recruitment despite occupying the same primary binding site [20–22]. Addressing these questions requires a dynamic perspective that captures allosteric coupling and conformational flexibility across receptor domains.

To address these gaps, we combine all-atom molecular dynamics (MD) simulations with multiscale CABS-flex modeling [23–25] to characterize how ligand binding redistributes receptor flexibility and controls access to signaling-competent states.

## 2.0 Methods

### 2.1 Molecular dynamics simulations

To complement the multiscale flexibility sampling obtained from CABS-flex, all-atom molecular dynamics (MD) simulations were performed to refine dynamic features and assess their stability in an explicit, atomistic environment. While CABS-flex efficiently explores large-scale conformational variability, MD provides time-resolved trajectories governed by explicit physical interactions, enabling validation and detailed characterization of functionally relevant motions.

MD simulations were carried out using Amber 22 [26] with the ff14SB force field [27]. A comprehensive set of systems covering the full functional cycle of the IL1 receptor was simulated independently. These included unbound components: interleukin-1 receptor type 1 (IL1R1; R1), interleukin-1 receptor antagonist (IL1Ra; Ra), and interleukin-1 (IL1), as well as antagonist- and agonist-bound receptor complexes (R1–Ra, R1–IL1), the fully assembled ternary signaling complex (R1–IL1–RAcP), and the R1–IL1 subcomplex extracted from the ternary structure (4DEP_binary). All simulations were initiated from experimentally determined receptor–ligand complexes to ensure that the analyzed interfaces correspond to biologically validated binding modes. To probe the intrinsic stability of ligand-induced conformations, additional simulations were performed in which ligands or receptors were removed from their respective complexes (denoted with the _apo or _Apo suffix). Short system identifiers corresponding to the original PDB entries are used throughout the manuscript and figure legends (system details are provided in Supplementary Table S1).

All systems were prepared, equilibrated, and simulated using an identical protocol. Detailed simulation parameters, system preparation steps, and equilibration procedures are provided in the Supplementary Methods. Production simulations were performed under NPT conditions at physiological temperature. To ensure sufficient sampling of key functional states, the unbound receptor (1G0Y) and antagonist-bound complex (1IRA) were simulated for 2.2 µs each, while all other systems were simulated for 1.0 µs. Trajectories were saved at regular intervals for subsequent analysis.

### 2.2 CABS-flex modeling

To complement all-atom MD simulations, we employed CABS-flex 3.0 [23], a well-established multiscale method for efficient simulation of protein structural flexibility [28–30]. By using a reduced representation and Monte Carlo sampling scheme, CABS-flex enables rapid exploration of near-native conformational dynamics at substantially lower computational cost than all-atom MD, while reproducing experimentally observed flexibility patterns and MD-derived dynamic trends [24,25,31].

Built upon the extensively validated CABS coarse-grained model [32], CABS-flex has been shown to reliably capture large-scale domain motions and intrinsic flexibility across diverse protein systems and has become a widely adopted tool for studying structure–dynamics–function relationships [24]. Importantly, its integration with all-atom reconstruction enables direct comparison between multiscale flexibility sampling and atomistic MD results, providing an effective framework for cross-validation of dynamic trends.

In this study, CABS-flex simulations were performed for the unbound receptor (1G0Y), the unbound antagonist (1ILR), the antagonist-bound complex (1IRA), and the agonist-bound complex (1ITB). The same starting structures were used for both CABS-flex and MD simulations to ensure consistency. For each system, 50 independent simulations were performed using default parameters. Resulting per-residue fluctuation profiles (RMSF) and reconstructed atomistic models were directly compared with MD-derived dynamics to assess ligand-dependent flexibility patterns across receptor states.

Additional methodological details are provided in the Supplementary Methods.

### 2.3 Conformational free energy landscape

To characterize and compare the conformational states sampled by the R1 receptor under different binding conditions, we constructed projected conformational free energy landscapes based on the combined MD simulation data. These landscapes are used as comparative representations of sampled conformational space, rather than as absolute thermodynamic free energy profiles of the receptor.

All MD trajectories were processed by removing solvent and ion coordinates. Conformations from all simulated systems—including the unbound receptor (1G0Y), antagonist-bound (R1–Ra, 1IRA), agonist-bound (R1–IL1, 1ITB and 4DEP_binary), the ternary complex (R1–IL1–RAcP, 4DEP), and all corresponding ligand-removed (apo) states—were pooled into a single combined ensemble. This procedure ensures that all receptor conformations are projected onto a shared conformational landscape, enabling direct comparison between functional states.

The internal dynamics of the combined ensemble were described using pairwise Cα–Cα distances, calculated with cpptraj [33,34]. This feature representation captures large-scale domain rearrangements that dominate receptor function. Dimensionality reduction was performed using a two-step protocol: initial denoising by principal component analysis (PCA), followed by time-lagged independent component analysis (tICA) to identify the slowest collective motions [35].

A tICA lag time of 20 ns was used to balance separation of slow, functionally relevant motions from fast local fluctuations on the microsecond timescale of the simulations. The resulting independent components (tICs) were used to generate two-dimensional projections of the conformational landscape. Projected probability densities were converted to potential of mean force (PMF) surfaces via Boltzmann inversion at 298 K [36].

Additional technical details are provided in the Supplementary Methods.

### 2.4 Analysis of protein dynamics and inter-residue contacts

Protein dynamics were characterized using per-residue fluctuation profiles (RMSF) and inter-residue contact analyses derived independently from CABS-flex and all-atom MD simulations. CABS-flex 3.0 webserver outputs provided RMSF profiles, residue–residue contact frequency maps, and cluster-representative structural models for each system. These data were used directly for comparison with atomistic MD results.

For MD simulations, RMSF values were calculated for Cα atoms using cpptraj [33]. To enable direct comparison of inter-residue contact patterns between CABS-flex and MD, equivalent numbers of representative frames were analyzed for both methods. Contact frequency maps derived from MD trajectories were computed and compared with those obtained from CABS-flex, allowing assessment of the consistency of interfacial interactions and dynamic coupling across modeling resolutions.

Detailed definitions of residue contacts, frame selection procedures, and visualization methods are provided in the Supplementary Methods.

## 3.0 Results and Discussion

### 3.1 Intrinsic flexibility and binding-induced stabilization predispose partners for recognition

To understand whether the binding partners possess intrinsic flexibility that predisposes them for receptor recognition, we characterized the conformational dynamics of both the antagonist (Ra) and the agonist (IL1) in their unbound and receptor-bound states using both all-atom molecular dynamics (MD) and CABS-flex simulations (Figure 2). Both proteins exhibit a consistent dynamic signature characteristic of the β-trefoil fold. This fold is defined by a structurally conserved and rigid hydrophobic core, exemplified in the antagonist Ra by residues 21–24, 44–47, and 66–69, which display low RMSF values. In contrast to this stable core, the surface-exposed loops and terminal segments of both proteins exhibit substantially higher mobility. These flexible interface loops may be subject to additional regulation by post-translational modifications, such as N-glycosylation [37].

**Figure 2.**
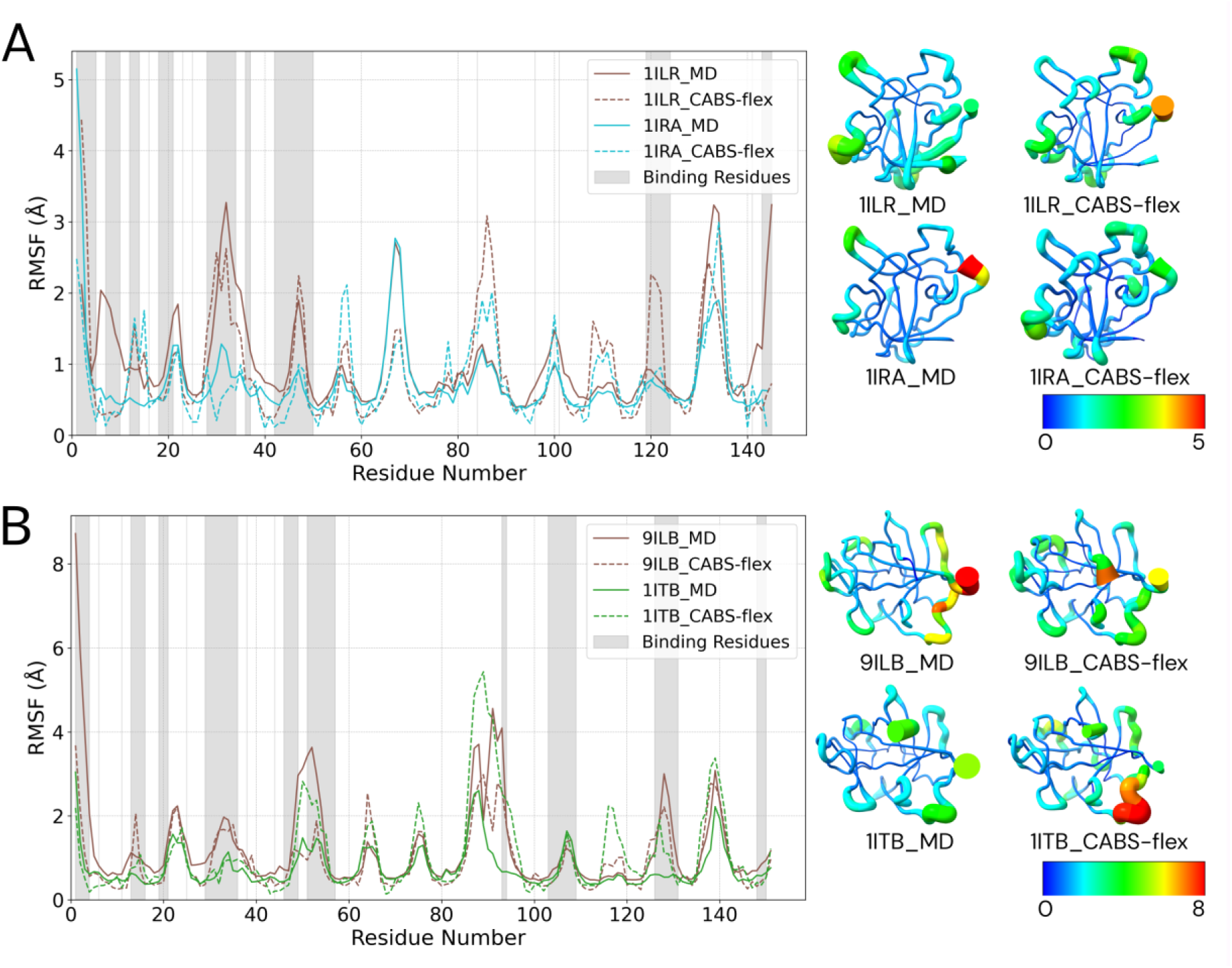
Intrinsic Flexibility and Binding-Induced Stabilization of the Agonist IL1 and Antagonist Ra. The RMSF profiles from all-atom MD (solid lines) and coarse-grained CABS-flex (dashed lines) simulations are compared for both the antagonist and agonist proteins in their unbound and receptor-bound states. Interface residues are highlighted with grey shading. **(A)** The antagonist Ra is more flexible in its unbound state (1ILR) than when complexed with the receptor (1IRA). **(B)** Similarly, the agonist IL1 is significantly more flexible in its unbound state (9ILB) compared to its receptor-bound form (R1–IL1, 1ITB). For both proteins, the binding-induced stabilization is most pronounced in the flexible loops that form the primary binding interface, demonstrating a shared mechanism of recognition and complex formation.

The RMSF profiles reveal a shared principle of binding-induced stabilization. The unbound antagonist Ra (1ILR) shows pronounced flexibility in its interface loops, which become significantly more rigid upon formation of the receptor complex (R1–Ra, 1IRA) (Figure 2A). A similar, and even more pronounced, stabilization is observed for the agonist IL1 (Figure 2B). The unbound agonist (9ILB) is highly dynamic, particularly in the loops that engage the receptor, and these same regions become markedly more rigid and ordered upon binding to R1 (R1-IL1, 1ITB).

Crucially, these regions of pronounced flexibility in the unbound proteins coincide with those previously implicated in receptor engagement. For example, the loop spanning residues 51–54 in Ra, which appears partially disordered in the unbound state, corresponds to a segment that interacts with the receptor’s D3 domain upon complex formation [13]. This shared mechanism—where a stable core preserves the binding-competent fold while localized flexibility in key loops allows for adaptive rearrangements before becoming ordered upon binding— highlights a balance between global structural stability and local conformational adaptability that facilitates efficient receptor recognition.

### 3.2 A conserved primary interface and divergent D3 engagement distinguish antagonist from agonist

To understand the molecular basis of competitive antagonism, we first compared the interaction interfaces of the antagonist-bound (1IRA) and agonist-bound (1ITB, 4DEP) complexes. Analysis of the crystal structures reveals that both the antagonist Ra and the agonist IL1 engage a remarkably similar footprint on the D1/D2 domains of the R1 receptor. This high degree of structural conservation at the primary binding site, evident from the overlap in interface residues, explains why the antagonist binds with high affinity and effectively competes with the agonist. This interface may be amenable to further optimization for higher affinity, for example through modulation of residues such as E127 that have been implicated in electrostatic complementarity [38].

Our inter-protein contact analysis of the simulated R1–Ra complex aligns perfectly with this observation (Figure 3). Both MD and CABS-flex simulations consistently showed stable interfacial contacts involving Tyr34, Gln36, and Tyr147, underscoring their central role in maintaining the primary receptor–ligand interface. This aligns closely with known receptor-binding determinants reported in biochemical and crystallographic studies. Notably, mutational studies of Tyr34, Gln36, and Tyr147 have shown reduced receptor affinity and interface stability, directly corroborating the contact patterns observed in our simulations [13,39]. Furthermore, our simulations captured high contact frequencies for Gln36 and its neighboring residue Asn39 [39]. This observation agrees with crystallographic data indicating that these residues form a stabilizing double hydrogen bond with the receptor backbone, contributing substantially to the overall binding energy. Together, these observations reinforce the robustness of the simulated binding pose.

**Figure 3.**
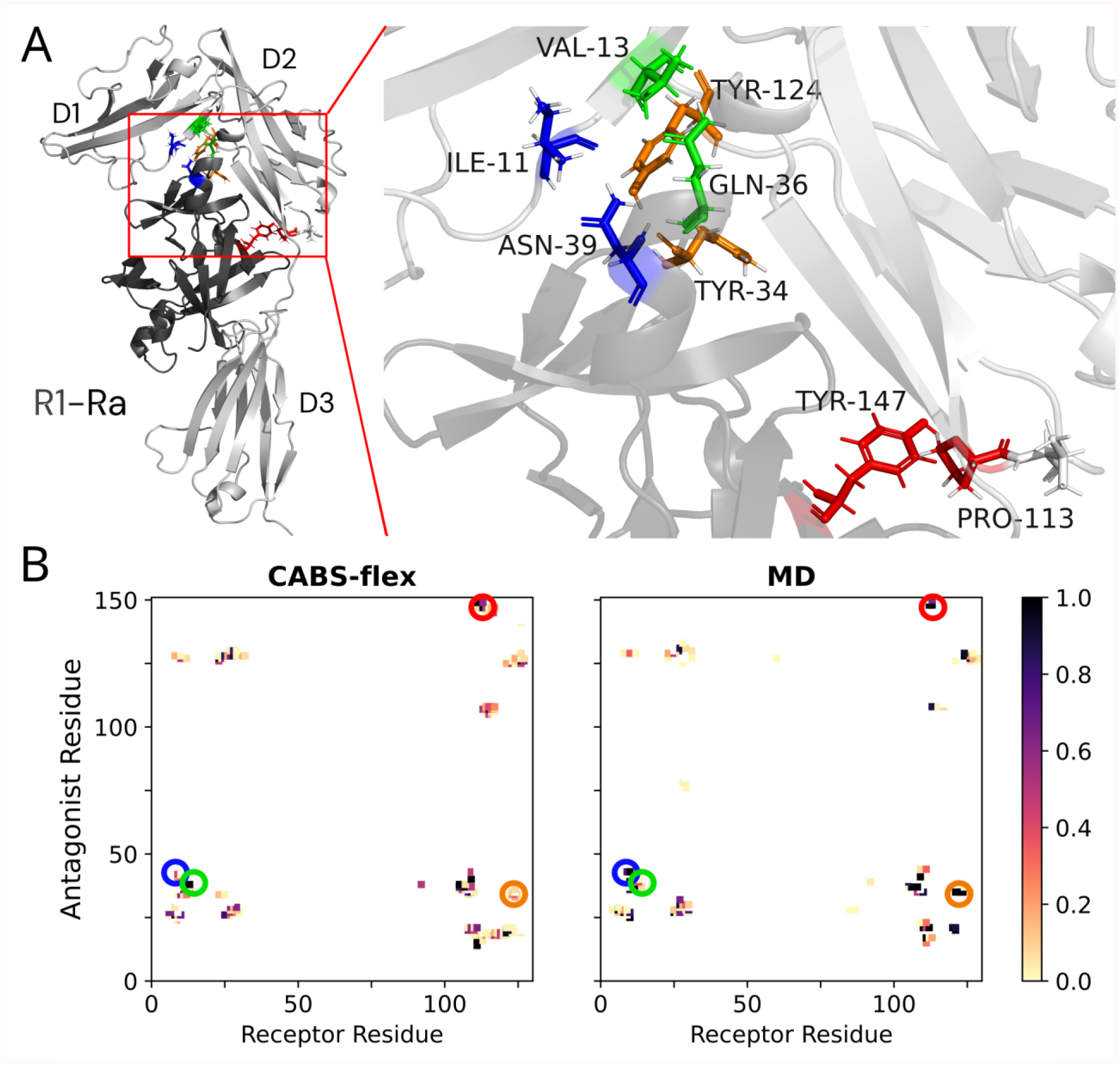
Key interfacial contacts mediating antagonist-receptor recognition. **(A)** Structure of the R1-Ra complex with key experimentally verified interface residues highlighted. Ra is shown in dark grey, and R1 in light grey. **(B)** Contact frequency maps from CABS-flex (left) and MD (right) simulations. High-frequency contacts (darker squares) from both methods align well with experimentally important residues (highlighted by colored circles), validating the simulated binding pose.

However, a crucial structural difference emerges at the D3 domain. The agonist acts as a "molecular clamp," forming an extensive secondary interface that functionally bridges the D1/D2 domains with D3 (Figure 1C). This locks all three domains into a rigid, signaling-competent architecture required to form the composite binding site for the co-receptor IL1-RAcP (Figure 1D).

In stark contrast, the antagonist lacks these key stabilizing contacts with D3 (Figure 1B). While it successfully occupies the primary site, it fails to tether the D3 domain. This structural difference—the antagonist’s failure to engage D3—is the origin of its opposing function. It suggests that the D3 domain becomes dynamically decoupled from the rest of the antagonist-bound receptor. This leads to the central hypothesis of our work: the lack of a stabilizing D3 interface effectively "frees" this domain, leading to an increase in its flexibility. We investigate this dynamic consequence in the following section.

### 3.3 Opposing dynamic signatures underlie allosteric discrimination by the IL1 receptor

The unbound R1 receptor is intrinsically dynamic and samples a broad conformational ensemble that is essential for its function. Analysis of the unbound receptor trajectories revealed substantial conformational heterogeneity, with interconversion between compact and extended states. This behavior is consistent with experimental SAXS and NMR studies on the IL1 receptor family, which demonstrated a pre-existing open–closed equilibrium in the absence of ligand [21,40]. Prior computational work has further shown that this intrinsic plasticity is largely driven by large-amplitude rotational freedom of the D3 domain in R1 and related receptors such as ST2 [22,41]. The D2–D3 interdomain linker residues have been identified as the primary hinge for these rotational movements[42].

A direct comparison of all-atom MD and CABS-flex simulations reveals strong qualitative agreement across all receptor states (Supplementary Figure S1), validating our multiscale approach. The conformational ensemble sampled by the unbound receptor (1G0Y) therefore defines a dynamic baseline against which the effects of agonist and antagonist binding can be evaluated (Figure 4A).

**Figure 4.**
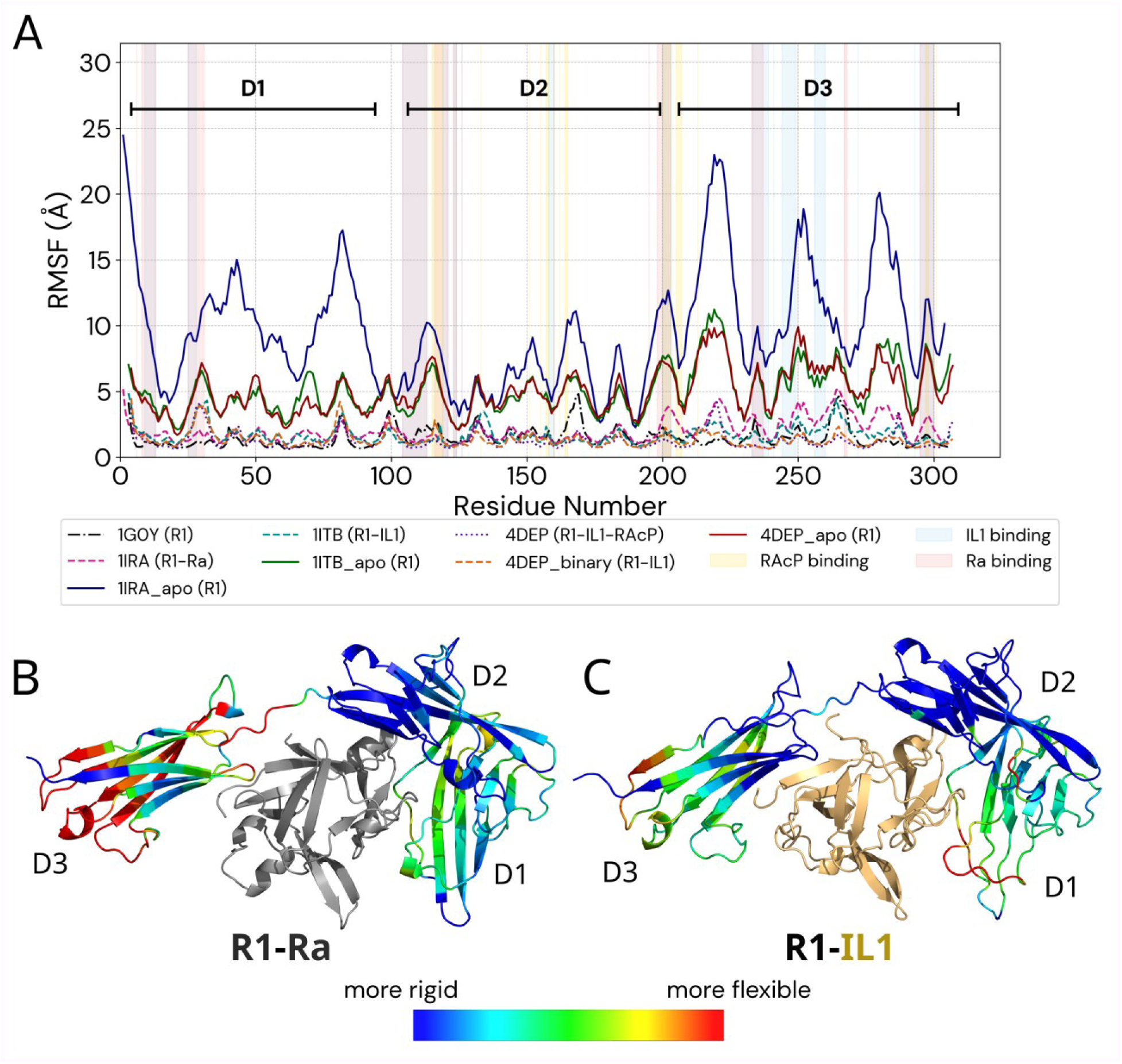
Opposing dynamic signatures underlie allosteric discrimination by the IL1 receptor. (A) RMSF profiles of the **R1** receptor reveal distinct dynamic pathways of inhibition and activation. The unbound receptor (1G0Y) defines a baseline of intrinsic flexibility. Antagonist binding (**R1–Ra**, 1IRA) uniquely increases the flexibility of the **D3** domain, particularly at the **RAcP** binding sites (shaded yellow). In contrast, agonist binding (**R1–IL1**, 1ITB) and formation of the ternary complex (**R1–IL1–RAcP**, 4DEP) lead to progressive stabilization. The pronounced hyper-flexibility of the receptor freed from the antagonist (1IRA_apo) reveals a latent instability induced by antagonist binding. (B, C) Changes in flexibility (dRMSF) relative to the unbound state are mapped onto the corresponding receptor structures, with dRMSF ≤ 0 shown in blue and dRMSF ≥ 1.5 Å in red. The maps highlight (B) rigidification of the primary antagonist-binding site coupled with increased flexibility of the distal D3 domain, versus (C) global stabilization and rigidification of the receptor upon agonist binding.

Antagonist binding disrupts this baseline in a highly specific manner. As shown in Figure 4, formation of the R1–Ra complex (1IRA) leads to a pronounced increase in flexibility of the distal D3 domain, while the D1/D2 domains remain comparatively rigid. This redistribution of flexibility is the direct dynamic consequence of the structural features identified above: the antagonist occupies the conserved primary binding site but fails to establish stabilizing contacts with D3. While crystallographic structures capture this lack of interdomain contacts [13–15], our simulations reveal its functional consequence—enhanced dynamic disorder of D3. This observation is further supported by elevated crystallographic B-factors in the D3 region of the R1–Ra complex.

Importantly, the antagonist-induced increase in flexibility is not uniform. Peaks in mobility are localized precisely at residues within D3 that form the binding interface for the co-receptor RAcP (Figure 4A). Specifically, these peaks in mobility correspond to critical residues such as Arg208 and Pro206, which are essential for docking the co-receptor. In the agonist-bound state, these residues are stabilized to form a ’polar hub’ and a hydrophobic patch with RAcP residues Tyr249 and Asn219, respectively (Supplementary Figure S2) [43]. By rendering this interface conformationally unstable, the antagonist effectively eliminates the structural platform required for stable RAcP association, thereby preventing signal transduction.

In stark contrast, activation proceeds through progressive stabilization of the receptor. Initial agonist binding (R1–IL1, 1ITB and 4DEP_binary) reduces receptor flexibility below the unbound baseline (Figure 4A). This rigidification provides a direct, atomistic explanation for early thermodynamic measurements reporting a substantial entropic penalty upon agonist binding, attributed to a restriction of receptor conformational freedom [40]. Full stabilization is achieved only upon formation of the ternary signaling complex (R1–IL1–RAcP, 4DEP), which represents the most rigid and conformationally stable state sampled. Activation thus corresponds to the stepwise construction of a rigid signaling platform. Complementary analysis of the agonist and antagonist proteins themselves confirms that their β-trefoil cores remain structurally stable across all states, while interface loops undergo consistent rigidification upon binding to R1 (Supplementary Figure S3).

The disruptive effect of the antagonist is further revealed by simulations of ligand-removed (“freed”) receptor states. RMSD profiles (Supplementary Figure S4) show that while bound complexes remain structurally stable, the receptor rapidly explores a vast conformational space upon ligand removal (exhibiting RMSD values up to 20 Å). When Ra is removed from the R1–Ra complex (1IRA_apo), the receptor rapidly explores a broad conformational space, with the D3 domain exhibiting large-amplitude fluctuations exceeding 20 Å. In contrast, receptors freed from agonist-bound states (R1–IL1 or R1–IL1–RAcP) display significantly attenuated D3 mobility. These results demonstrate that antagonist binding does not simply fail to stabilize the receptor, but actively forces it into a strained, sparsely sampled conformation that is intrinsically unstable once the antagonist is removed.

Collectively, these opposing dynamic signatures provide a clear mechanistic explanation for allosteric discrimination by R1. Activation is driven by progressive stabilization of receptor dynamics, whereas antagonism operates through targeted dynamic disruption. By increasing conformational variability at the critical D3 interface, antagonist binding makes formation of the stable RAcP-recruiting platform physically inaccessible, thereby locking the receptor in a signaling-incompetent state.

### 3.4 The conformational landscape defines the allosteric pathways of inhibition and activation

To place the observed dynamic signatures into a unified mechanistic framework, we mapped the conformational energy landscape of the R1 receptor (Figure 5). This analysis allows visualization of how different ligands bias the receptor toward distinct, low-energy conformational states and reveals that activation and inhibition proceed along separate and mutually exclusive allosteric pathways.

**Figure 5.**
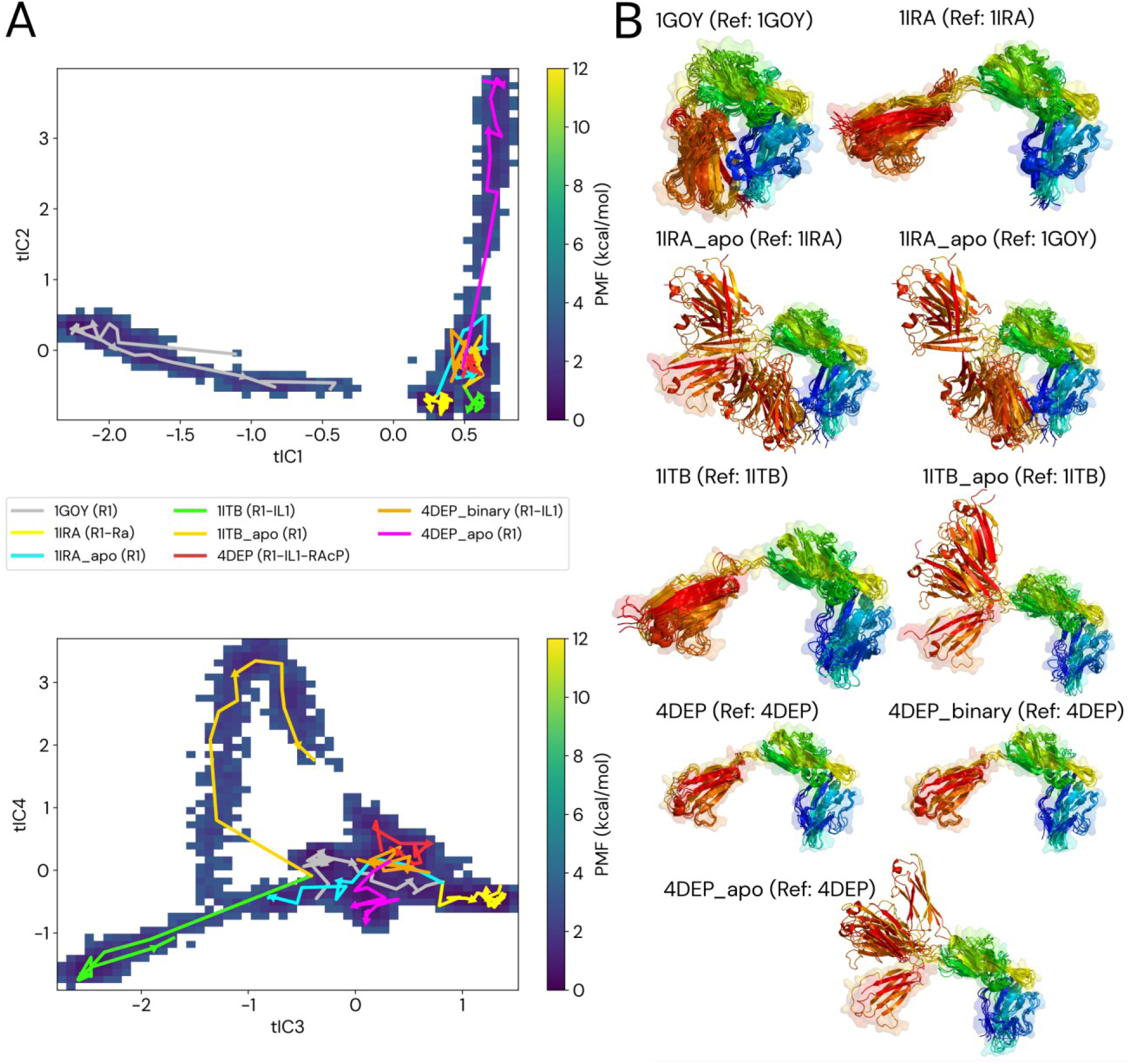
Conformational landscape defines the allosteric pathways of inhibition and Activation. (A) The potential of mean force (PMF) landscape of the R1 receptor projected onto two complementary sets of time-independent components (tICs). Dark blue regions indicate low-energy conformational basins. Overlaid trajectories illustrate the behavior of key functional states along these collective coordinates. (B) Structural ensembles corresponding to each simulation are shown, with frames sampled at 200 ns intervals and colored by residue index (N-terminus blue, C-terminus red). The initial reference structure is shown as a semi-transparent surface. The projections reveal that the antagonist-bound receptor (R1–Ra, 1IRA) is confined to a distinct low-energy basin corresponding to the inhibited conformation. In contrast, the unbound receptor (1G0Y) and ligand-removed (“apo”) simulations display extensive exploratory behavior. Agonist-bound binary complexes (R1–IL1, 1ITB and 4DEP_binary) populate a separate conformational basin that represents an intermediate along the activation pathway, while the fully assembled ternary complex (R1–IL1–RAcP, 4DEP) stabilizes the fully active state.

Activation follows a stepwise pathway guided by the agonist. In the tIC1–tIC2 projection (Figure 5A), agonist-bound binary complexes (R1–IL1, 1ITB and 4DEP_binary) induce a clear population shift from the inhibited basin, populated by the unbound receptor (1G0Y), into a distinct intermediate basin. Structural ensembles sampled along this transition reveal a coordinated, large-scale reorientation of the receptor domains (Figure 5B), which is required before co-receptor recruitment can occur. Full stabilization of the active state is achieved only upon formation of the complete ternary complex (R1–IL1–RAcP, 4DEP), which occupies a well-defined, low-energy basin corresponding to the signaling-competent conformation. This separation of inactive, intermediate, and active basins is robust across alternative projections of the conformational landscape (Supplementary Figure S5).

In contrast, antagonism operates through energetic trapping. In the tIC3–tIC4 projection (Figure 5A), the antagonist-bound receptor (R1–Ra, 1IRA) is confined to the inhibited basin, despite the intrinsic flexibility of the unbound receptor, which explores a broad conformational space. This confinement reflects the energetic consequence of the antagonist-induced dynamic disorder in the D3 domain described in the previous section (Figure 4). Rather than guiding the receptor toward an activation pathway, antagonist binding restricts access to alternative conformational states required for signaling.

Simulations of ligand-removed (“apo”) states further clarify the nature of this trapping. When Ra is removed, the receptor (1IRA_apo) rapidly escapes the inhibited basin and explores a wide region of conformational space, relaxing toward the ensemble characteristic of the unbound receptor (1G0Y). Structural ensembles in Figure 5B illustrate this relaxation: relative to the antagonist-bound reference, the D3 domain undergoes large-amplitude reorientation, whereas superposition onto the unbound reference reveals convergence toward a less strained architecture. These observations confirm that antagonist binding stabilizes a high-energy, strained receptor conformation that is intrinsically unstable once the antagonist is removed.

### 3.5 An integrative model for the allosteric control of IL1 receptor signaling

Previous structural and biophysical studies have established the modular organization of the IL1 receptor and identified the D3 domain as a key determinant of signaling competence [17,21,22]. Recent molecular dynamics analyses of other IL-1 family cytokines, including IL-37, further emphasize the role of conformational stability and interface dynamics in regulating functional state transitions [44]. This view is reinforced by recent high-resolution cryo-EM structures of related receptors, including IL-36R, which reveal an intrinsically flexible D3 domain that becomes ordered only upon formation of the active ternary complex [18]. Notably, a recent structural study of a small-molecule IL1 antagonist (PDB 8C3U) [45] reports displacement of an R1-interacting loop in IL1; structural alignment with the IL1–R1 complex suggests that such rearrangement could interfere with productive D3 engagement, further underscoring the importance of correct D3 positioning for signaling competence.

Building on these observations, our results define a dynamic model that links ligand binding to receptor function through redistribution of conformational flexibility. Agonist binding promotes progressive stabilization of receptor dynamics, guiding R1 through a stepwise pathway toward formation of a rigid, signaling-competent platform. This stabilization culminates in recruitment of the co-receptor RAcP, which locks the receptor into its fully active conformation.

In contrast, antagonist binding at the conserved D1/D2 interface selectively increases conformational variability in the distal D3 domain. This targeted dynamic disruption destabilizes the co-receptor binding interface and prevents productive assembly of the signaling complex. Rather than acting through simple steric blockade, antagonism arises from allosteric reshaping of the receptor’s dynamic landscape. Consistent with this mechanism, specific IL1 residues involved in IL1-RAcP engagement (e.g., Gly140 and Gln141 in the ternary complex) are not fully conserved in IL-1Ra, further reducing the capacity of the antagonist-bound receptor to support stable co-receptor interactions [46].

Together, these observations support a population-shift mechanism in which agonists and antagonists bias the IL1 receptor toward distinct regions of conformational space. Stabilization of signaling-competent states underlies activation, whereas antagonist-induced disorder traps the receptor in a signaling-incompetent ensemble. These conclusions pertain to the extracellular ectodomain of IL1R1, which constitutes the ligand-recognition and co-receptor recruitment module. Integration with the transmembrane and intracellular regions may further shape the conformational landscape in the cellular context.

## 4.0 Conclusions

In this work, we show that discrimination between agonist and antagonist binding by the IL1 receptor is governed by opposing effects on receptor dynamics rather than by stabilization of alternative static structures. Ligands that bind to the same conserved site impose distinct dynamic signatures that determine signaling outcome.

Activation emerges as a process of progressive stabilization, in which agonist binding and subsequent co-receptor recruitment incrementally rigidify the receptor to form a stable signaling platform. Antagonism, by contrast, is an active process of dynamic disruption, in which increased flexibility of the D3 domain renders the co-receptor binding interface functionally inaccessible.

By mapping the receptor’s conformational landscape, we demonstrate how this antagonist-induced disorder energetically traps the receptor in a signaling-incompetent state and prevents entry into the activation pathway. This dynamic population-shift mechanism reconciles long-standing structural and functional observations and explains how ligands binding to the same site can produce opposite biological outcomes.

Beyond the IL1 system, our findings suggest a general principle for cytokine receptor regulation in which signaling outcomes are governed by ligand-induced redistribution of receptor flexibility rather than by stabilization of alternative static conformations. In multi-component receptor systems, distal regulatory domains that control co-receptor recruitment may therefore represent effective targets for allosteric modulation, offering an alternative to strategies focused solely on ligand sequestration or direct interface blockade.

## Data Availability

The molecular dynamics trajectories generated in this study are available in the Mendeley Data repository at https://doi.org/10.17632/sw7ppcknm8.

## Declaration of generative AI and AI-assisted technologies in the writing process

During the preparation of this work, the authors used ChatGPT (OpenAI), AI studio (Google) and Claude (Anthropic) to proof-read the text to enhance the quality of the language. After using these tools, the authors reviewed and edited the content as needed and took full responsibility for the content of the publication.

## Acknowledgements

S.K., A.F, and C.N. acknowledge funding from the National Science Centre, Poland (SHENG 2021/40/Q/NZ2/00078). We gratefully acknowledge Polish high-performance computing infrastructure PLGrid (HPC Center: ACK Cyfronet AGH) for providing computer facilities and support within computational grant nos. PLG/2025/017952 and PLG/2026/019105.

## CRediT author statement

**CN**: Conceptualization, Data curation, Formal analysis, Investigation, Methodology, Resources, Software, Supervision, Validation, Visualization, Writing – original draft, Writing – review & editing. **AF:** Data curation, Formal analysis, Investigation, Methodology, Software, Validation, Visualization, Writing – original draft, Writing – review & editing. **SK**: Conceptualization, Formal analysis, Funding acquisition, Investigation, Methodology, Project administration, Resources, Supervision, Validation, Visualization, Writing – original draft, Writing – review & editing.

## Supplementary Materials

Supplementary Methods 1. Detailed Molecular Dynamics Simulation Protocol

Supplementary Methods 2. CABS-flex simulations

Supplementary Methods 3. Conformational free energy landscape analysis

Supplementary Methods 4. Analysis of protein dynamics and inter-residue contacts

Supplementary Table S1. Summary of all molecular dynamics systems simulated in this study

Supplementary Figure S1. Comparative dynamic profiles of the R1 receptor from all-atom MD and coarse-grained CABS-flex simulations.

Supplementary Figure S2. Key interfacial contacts between the primary receptor (R1) and the co-receptor (RAcP).

Supplementary Figure S3. Intrinsic and bound state dynamics of the agonist IL1 and antagonist Ra.

Supplementary Figure S4. Backbone RMSD for all systems simulated with molecular dynamics.

Supplementary Figure S5. Alternative projections of the R1 conformational free energy landscape.

