## Supplementary for "Antagonist binding actively disrupts interleukin-1 receptor dynamics to block co-receptor recruitment"

### Supplementary Methods

#### 1. Detailed Molecular Dynamics Simulation Protocol

All-atom molecular dynamics (MD) simulations were performed using the Amber 22 software package [1] with the PMEMD.cuda engine [2–4]. Protein interactions were described using the AMBER ff14SB force field [5]. Initial coordinates were taken from experimentally determined structures deposited in the Protein Data Bank [6] (PDB): 1G0Y, 1ILR, 9ILB, 1IRA, 1ITB, and 4DEP. The R1–IL1 binary complex extracted from the ternary structure is referred to as 4DEP_binary. Detailed information on starting PDB entries and chain composition for all simulated systems is provided in Supplementary Table S1. Missing residues present in the crystallographic structures were modeled prior to simulation using SWISS-MODEL [7]. The modeled segments were subjected to the same minimization and equilibration protocol as the remainder of the system.

#### System preparation

Each system was prepared using the tleap module from AmberTools [8]. All crystallographic water molecules present in the original PDB structures were removed prior to system setup. Proteins were solvated in a truncated octahedral box of TIP3P water molecules [9] with a minimum buffer distance of 10.0 Å between the solute and the box boundary. Systems were neutralized and adjusted to physiological ionic strength by addition of $Na^+$ and $Cl^-$ ions using Joung–Cheatham parameters [10] optimized for the selected water model.

#### Energy minimization

Energy minimization was carried out in two stages, each consisting of 10,000 cycles. In the first stage, positional restraints were applied to all protein atoms with a force constant of 20.0 kcal $mol^{-1}$ $Å^{-2}$ to relax solvent and ions. In the second stage, all restraints were removed and the entire system was minimized.

#### Heating and equilibration

Systems were gradually heated from 100 K to 300 K over 500 ps under NVT conditions using a Langevin thermostat [11] with a collision frequency of 5.0 $ps^{-1}$. During heating, positional restraints of 20.0 kcal $mol^{-1}$ $Å^{-2}$ were maintained on protein atoms.

Subsequently, density equilibration was performed for 500 ps under NPT conditions at 300 K and 1 atm using a Monte Carlo barostat [12], while maintaining protein restraints. Positional restraints on backbone atoms (CA, C, N, O) were then gradually released over three 200 ps stages under NVT conditions, with restraint force constants reduced sequentially from 10.0, to 5.0, and finally 1.0 kcal $mol^{-1}$ $Å^{-2}$. A final 2.0 ns unrestrained equilibration under NVT conditions was performed to ensure full relaxation of the system.

**Production simulations**

Production MD simulations were carried out under NPT conditions at 298 K and 1 atm using a Langevin thermostat [11] with a collision frequency of 1.0 $ps^{-1}$ and isotropic pressure coupling. A 2.0 fs integration timestep was used, with bonds involving hydrogen atoms constrained using SHAKE [13] and rigid water geometry enforced using SETTLE [14]. Long-range electrostatic interactions were treated using the Particle Mesh Ewald (PME) method [15,16] with a real-space cutoff of 12.0 Å.

To ensure sufficient sampling of key functional states, the unbound receptor (1G0Y) and antagonist-bound complex (1IRA) were simulated for a total of 2.2 μs each, while all other systems were simulated for 1.0 μs each. Simulations were extended by restarting from previous coordinates and velocities where appropriate. System coordinates were saved every 1.0 ps for subsequent analysis.

Simulations were performed as continuous microsecond-scale trajectories rather than multiple short replicas in order to allow uninterrupted sampling of slow interdomain motions and large-scale conformational transitions relevant to receptor activation and inhibition. The primary objective was to characterize ligand-dependent redistribution of conformational populations along collective coordinates. System stability and convergence were assessed via backbone RMSD (Supplementary Figure S4), which confirmed that all bound and reference systems reached structural equilibrium.

### 2. CABS-flex simulations

CABS-flex simulations were performed using the CABS-flex 3.0 web server [17]. The method is based on the CABS coarse-grained protein model [18] and employs Replica Exchange Monte Carlo (REMC) sampling to efficiently explore near-native conformational flexibility. CABS-flex has been extensively benchmarked against experimental data and all-atom MD simulations and has been shown to accurately reproduce intrinsic protein flexibility and large-scale domain motions.

Each system was subjected to 50 independent REMC simulations using default simulation settings. Coarse-grained trajectories were clustered using the standard CABS-flex analysis pipeline. All-atom reconstructions were generated from representative coarse-grained models using server-provided reconstruction tools. Per-residue fluctuation profiles (RMSF) were extracted from the reconstructed ensembles and compared directly with RMSF profiles obtained from all-atom MD simulations.

### 3. Conformational free energy landscape analysis

To characterize the conformational space sampled by the R1 receptor across all simulated conditions, a combined-ensemble analysis was performed. All-atom MD trajectories were first processed by removing solvent and ion coordinates. Receptor conformations from all simulations—including the unbound receptor (1G0Y), antagonist-bound (R1–Ra, 1IRA), agonist-bound (R1–IL1, 1ITB and 4DEP_binary), the ternary signaling complex (R1–IL1–RAcP, 4DEP), and all corresponding ligand-removed (apo) states—were pooled into a single dataset. This approach ensures that all conformations are projected onto a common conformational space, enabling direct comparison between functional states. For consistency across systems, equivalent receptor residue ranges (residues 4–307) were used in the pooled ensemble employed for dimensionality reduction and free energy landscape construction.

**Feature selection and dimensionality reduction**

The internal dynamics of the combined ensemble were described using pairwise Cα–Cα distances calculated with cpptraj [19] from the AmberTools package [8]. This feature representation captures large-scale domain rearrangements and interdomain motions that dominate the functional behavior of the receptor, while remaining insensitive to high-frequency local fluctuations.

To reduce dimensionality and extract the slow collective motions governing receptor dynamics, a two-step dimensionality reduction protocol was applied. First, principal component analysis (PCA) was performed using IncrementalPCA from the scikit-learn package [20] to remove high-dimensional noise and reduce correlations in the feature space. The PCA-transformed data were subsequently analyzed using time-lagged independent component analysis (tICA) using the deeptime library [21], which identifies collective variables associated with the slowest decorrelating motions in the system [22]. All numerical processing and data manipulation were handled using python modules NumPy [23] and Pandas [24], with final visualizations generated via Matplotlib [25].

A tICA lag time of 20 ns was selected based on the characteristic timescales observed in the MD trajectories, representing a compromise between filtering fast local motions and retaining sufficient statistical sampling of slow transitions on the microsecond timescale. The top **ten** independent components were retained for subsequent analysis. Alternative lag times and component numbers were tested and yielded qualitatively similar conformational projections (data not shown).

**Construction of free energy landscapes**

Two-dimensional conformational landscapes were generated by projecting the combined-ensemble trajectories onto pairs of selected tICs. The probability density of sampled conformations in each projection was estimated using histogram-based methods [26]. The corresponding potential of mean force (PMF) surfaces were calculated via Boltzmann inversion according to:

$$PMF(x, y) = -k_B T \ln P(x, y)$$

where $P(x, y)$ is the normalized probability density, $k_B$ is the Boltzmann constant, and $T$ is temperature ( $T = 298\ K$).

The resulting PMF surfaces are interpreted as comparative representations of the sampled conformational space, rather than as absolute thermodynamic free energy profiles. Their primary purpose is to visualize population shifts, energetic basins, and transition pathways associated with different functional states of the receptor.

### 4. Analysis of protein dynamics and inter-residue contacts

Per-residue fluctuation profiles (RMSF) were obtained directly from the CABS-flex 3.0 webserver outputs for each simulated system. The webserver provides RMSF profiles calculated from ensembles of 1,000 coarse-grained models per system, as well as residue–residue contact frequency maps and cluster-representative structures.

For all-atom MD simulations, RMSF values were calculated for Cα atoms using cpptraj [19] from the AmberTools package [8]. Trajectories were aligned to a common reference structure prior to RMSF calculation to remove overall translational and rotational motion.

To enable direct comparison of contact frequencies between CABS-flex and MD simulations, MD trajectories were processed using the MDAnalysis Python package [27]. From each MD trajectory, 1,000 representative frames were extracted to match the sampling density of the CABS-flex ensembles. Residue–residue contacts were defined based on the distance between side-chain centers of mass, with a distance cutoff of 6.8 Å used to define contact formation.

Contact frequencies were calculated as the fraction of analyzed frames in which a given residue pair satisfied the contact criterion. The resulting contact frequency matrices were visualized as heatmaps using python modules Matplotlib [25] and Seaborn [28].

**Supplementary Table S1. Summary of all molecular dynamics systems simulated in this study.**

| System label | Starting PDB | Chains used | Description |
|---|---|---|---|
| R1 | 1G0Y | R | Unbound receptor |
| Ra | 1ILR | 1 | Unbound antagonist |
| IL1 | 9ILB | A | Unbound agonist |
| R1–Ra | 1IRA | Y (R1), X (Ra) | Antagonist-bound complex |
| R1–IL1 | 1ITB | B (R1), A (IL1) | Agonist-bound complex |
| R1–IL1–RAcP | 4DEP | B (R1), A (IL1),<br><br>C (RAcP) | Fully assembled ternary complex |
| 4DEP_binary | 4DEP | B (R1), A (IL1) | R1–IL1 binary extracted from ternary |
| 1IRA_apo | 1IRA | Y (R1 only) | Receptor freed from Ra |
| 1ITB_apo | 1ITB | B (R1 only) | Receptor freed from IL1 |
| 4DEP_apo | 4DEP | B (R1 only) | Receptor freed from ternary |
| 1IRA_Apo | 1IRA | X (Ra only) | Antagonist freed from R1 |
| 1ITB_Apo | 1ITB | A (IL1 only) | Agonist freed from R1 |
| 4DEP_Apo | 4DEP | A (IL1 only) | Agonist freed from ternary |

### Supplementary Figures

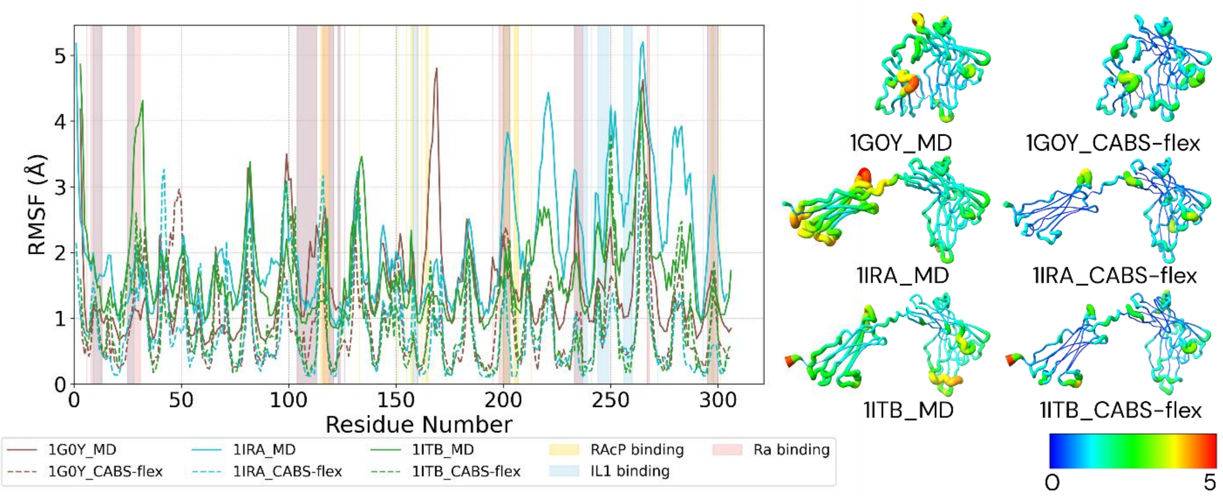


**Supplementary Figure S1. Comparative dynamic profiles of the R1 receptor from all-atom MD and coarse-grained CABS-flex simulations.** The figure compares the per-residue root-mean-square fluctuation (RMSF) for the R1 receptor in its unbound (1G0Y), antagonist-bound (1IRA), and agonist-bound (1ITB) states. The RMSF plot (left) and corresponding 3D structural mappings (right) show strong qualitative agreement between the dynamic hotspots identified by MD (solid lines) and CABS-flex (dashed lines), validating the multi-scale approach. The comparison also visually confirms the opposing dynamic principles of inhibition versus activation: antagonist binding increases the flexibility of the D3 domain relative to the unbound state, particularly at the RAcP binding sites (yellow highlights), while agonist binding leads to a general stabilization of the receptor.

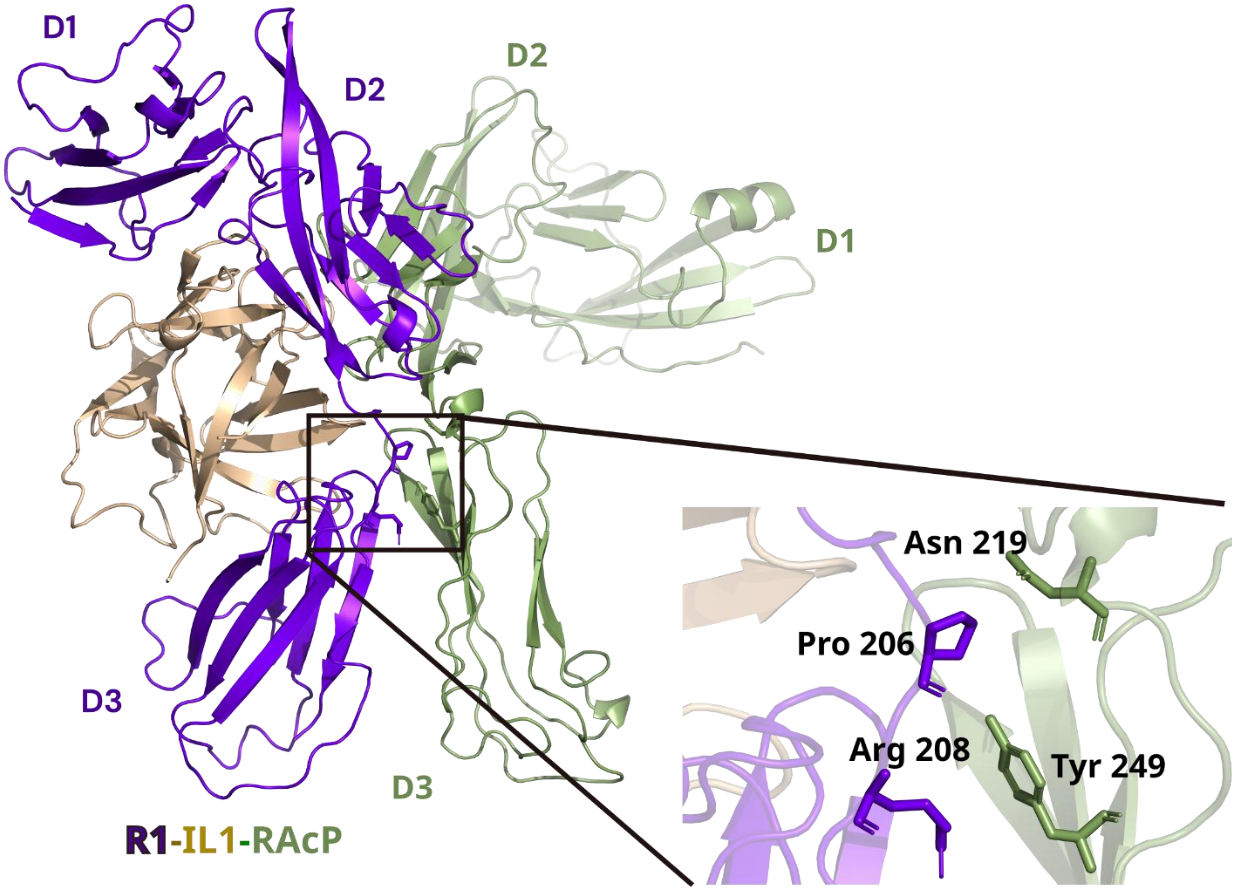


**Supplementary Figure S2. Key interfacial contacts between the primary receptor (R1) and the co-receptor (RAcP).** Structural representation of the active ternary signaling complex (PDB: 4DEP). The primary receptor (R1) is shown in purple, the agonist (IL-1) in gold, and the co-receptor (RAcP) in olive green. The inset provides a detailed view of the secondary binding interface. Key residues involved in recruitment include Pro206 and Arg208 on the R1-D3 domain, which form critical contacts with Asn219 and Tyr249 on RAcP. The increased flexibility of these R1 residues in the antagonist-bound state (as shown in Figure 4A) prevents the formation of this stable interface, thereby blocking co-receptor recruitment.

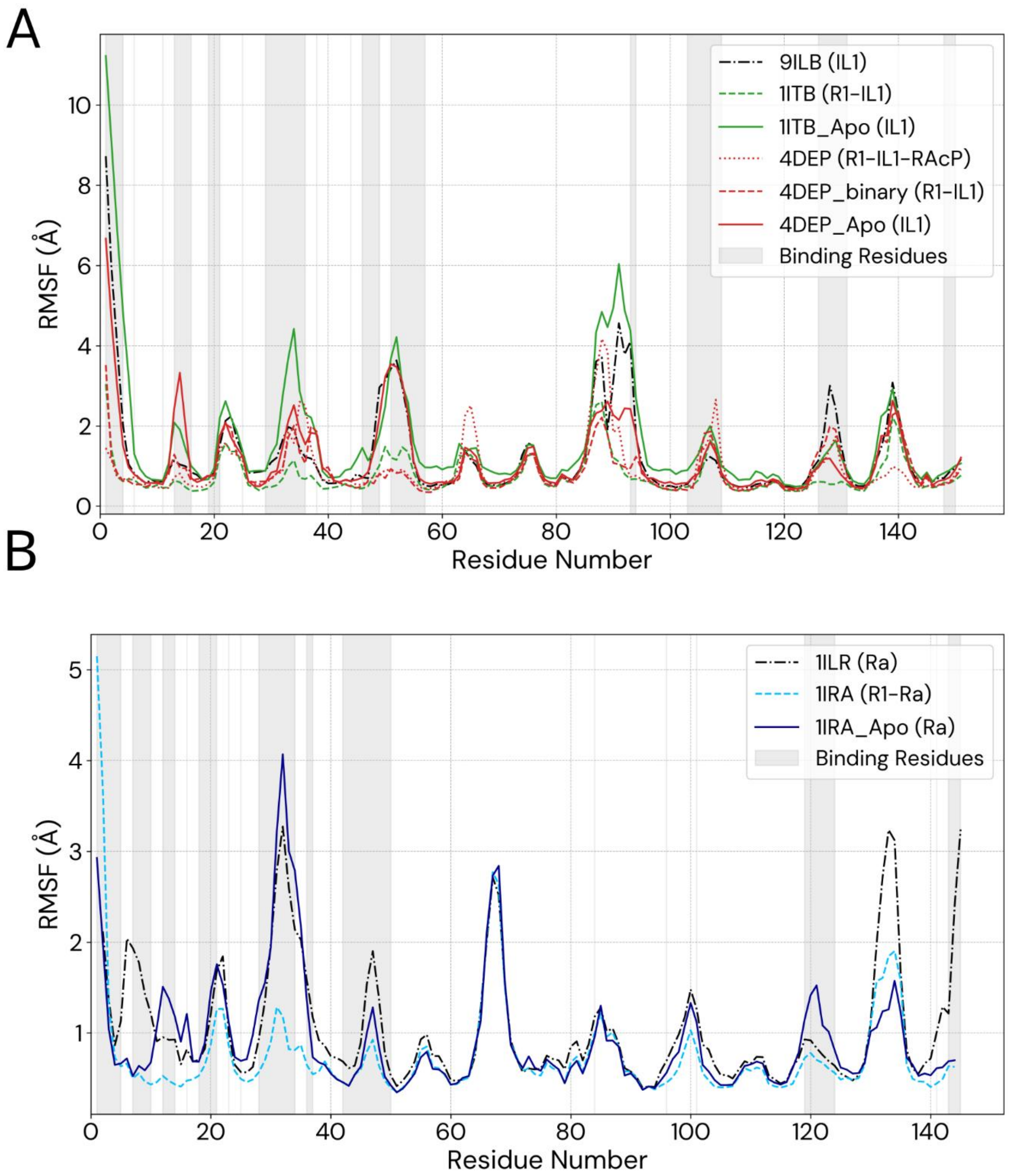


**Supplementary Figure S3. Intrinsic and bound state dynamics of the agonist IL1 and antagonist Ra.** The RMSF profiles for the agonist and antagonist proteins are shown across their key simulated states. Residues that form the primary binding interface with the R1 receptor are highlighted in grey. (A) The agonist cytokine IL1 is shown in its unbound state (from PDB: 9ILB), bound to R1 in the binary complex (1ITB), bound to R1 in the ternary complex (4DEP, 4DEP_binary), and its corresponding "freed" state (1ITB_Apo, 4DEP_Apo). (B) The antagonist protein Ra is shown in its unbound state (from PDB: 1ILR), bound to R1 (1IRA), and its corresponding "freed" state (1IRA_Apo). All proteins exhibit a dynamic signature characteristic of the β-trefoil fold, with a stable core and higher flexibility in the termini and surface-exposed loops.

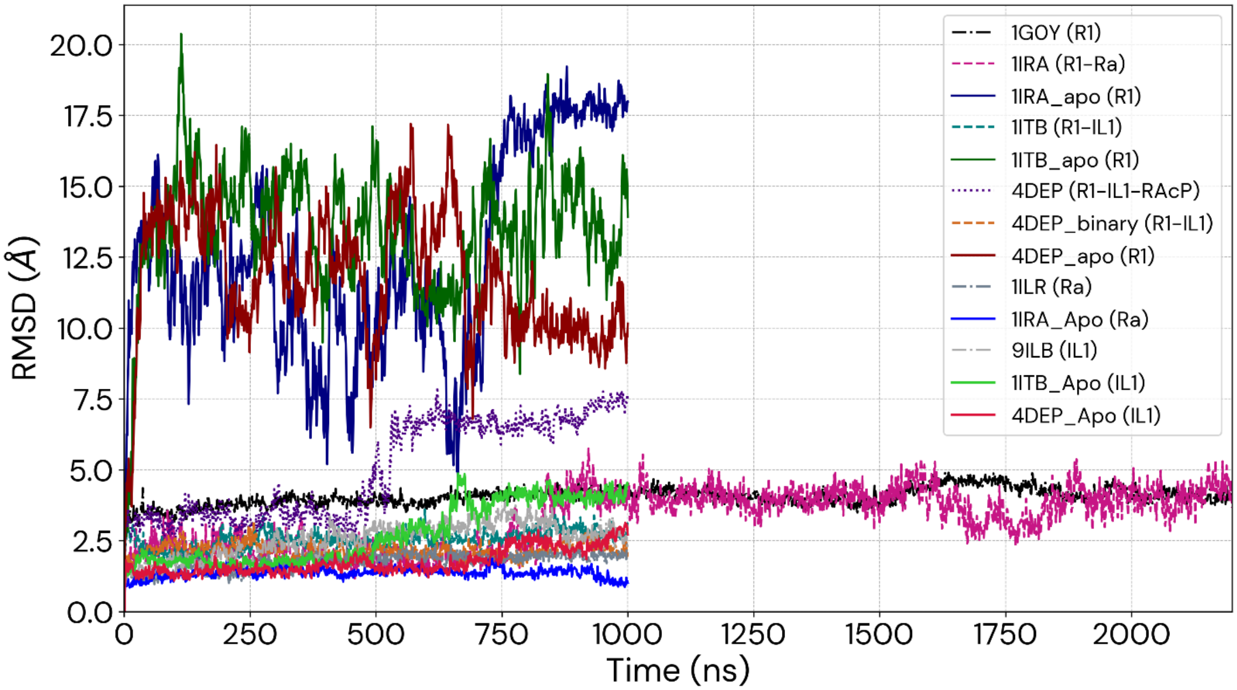


**Supplementary Figure S4. Backbone RMSD for all systems simulated with molecular dynamics.** Backbone RMSD (calculated for CA, C, and N atoms) is shown relative to the initial experimental or modeled starting structure. The unbound references (1G0Y, 1ILR, 9ILB) and bound complexes (1IRA, 1ITB, 4DEP) exhibit structural convergence and stability throughout the 1.0–2.2 μs production trajectories. The ligand-removed ("freed") receptor states, denoted with the *_apo* suffix, show significantly higher RMSD values (10–20 Å). This reflects the extensive conformational exploration and large-amplitude reorientation of the D3 domain that occurs once the stabilizing or straining effect of the ligand is removed, as described in the main text. Note that the 1G0Y (unbound R1) and 1IRA (antagonist-bound complex) were simulated for an extended duration of 2.2 μs to ensure thorough sampling of these critical states.

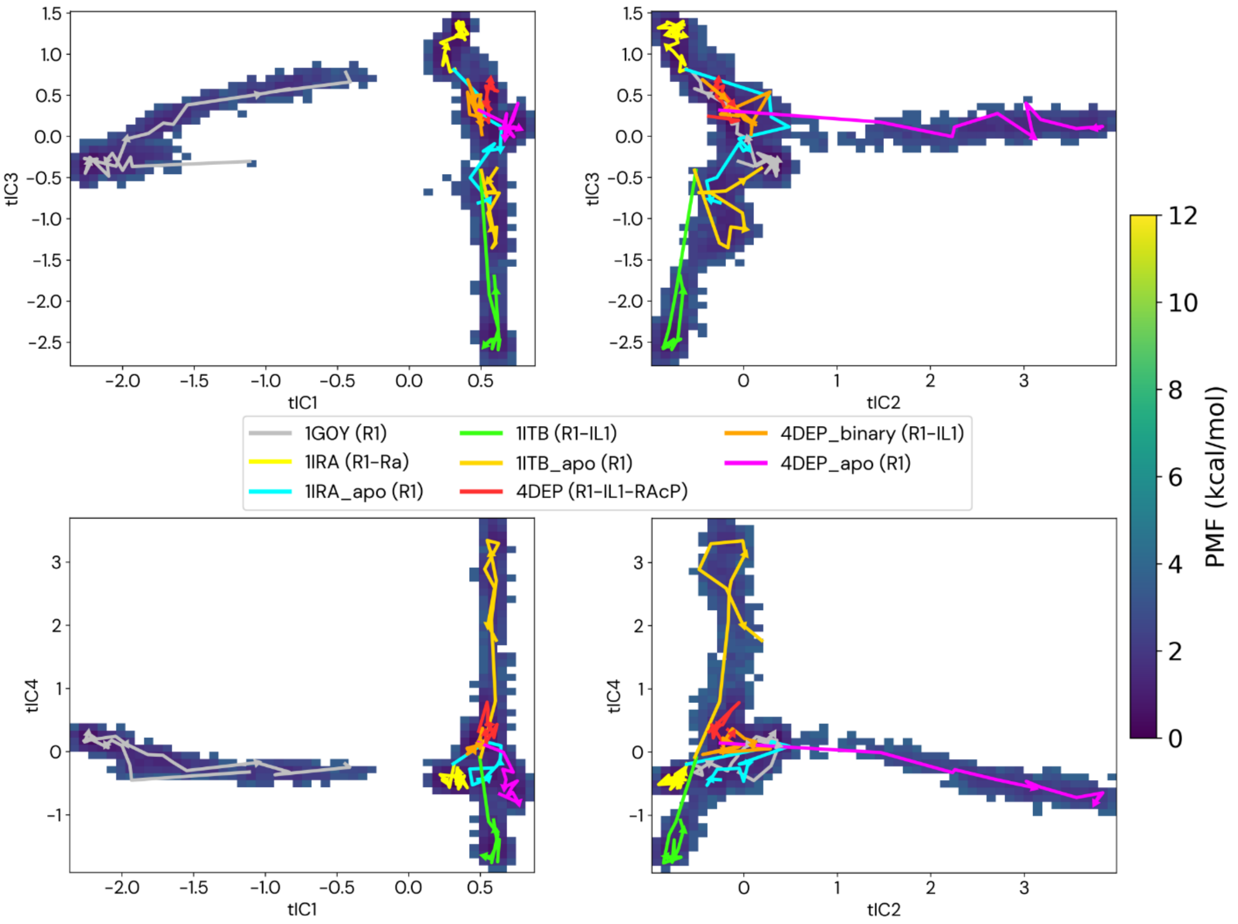


**Supplementary Figure S5. Alternative projections of the R1 conformational free energy landscape.** The Potential of Mean Force (PMF) landscape of the R1 receptor is projected onto four different pairs of time-independent components (tICs) to illustrate the robustness of the conformational analysis. Across all projections, the trajectories for the key functional states occupy distinct regions of the landscape. The antagonist-bound state (1IRA) is consistently confined to the same low-energy basin as the unbound receptor (1G0Y), while the agonist-bound complexes (1ITB, 4DEP_binary, 4DEP) populate a separate, distinct conformational region. This consistency across multiple views confirms the central findings that the antagonist confines the receptor to a restricted region of the conformational landscape in an inhibited state, preventing its transition to the agonist-bound intermediate state required for activation.

[8] D.A. Case, H.M. Aktulga, K. Belfon, D.S. Cerutti, G.A. Cisneros, V.W.D. Cruzeiro, N. Forouzesh, T.J. Giese, A.W. Götz, H. Gohlke, S. Izadi, K. Kasavajhala, M.C. Kaymak, E. King, T. Kurtzman, T.S. Lee, P. Li, J. Liu, T. Luchko, R. Luo, M. Manathunga, M.R. Machado, H.M. Nguyen, K.A. O'Hearn, A. V. Onufriev, F. Pan, S. Pantano, R. Qi, A. Rahnamoun, A. Risheh, S. Schott-Verdugo, A. Shajan, J. Swails, J. Wang, H. Wei, X. Wu, Y. Wu, S. Zhang, S. Zhao, Q. Zhu, T.E. Cheatham, D.R. Roe, A. Roitberg, C. Simmerling, D.M. York, M.C. Nagan, K.M.

Merz, AmberTools, J. Chem. Inf. Model. 63 (2023) 6183–6191. https://doi.org/10.1021/ACS.JCIM.3C01153.
